# Dynamic Modeling of Signal Transduction by mTOR Complexes in Cancer

**DOI:** 10.1101/633891

**Authors:** Mohammadreza Dorvash, Mohammad Farahmandnia, Pouria Mosaddeghi, Mitra Farahmandnejad, Hosein Saber, Mohammadhossein Khorraminejad-Shirazi, Amir Azadi, Iman Tavassoly

## Abstract

Signal integration in the mTOR pathway plays a vital role in cell fate decision making in cancer cells. As a signal integrator, mTOR shows a complex dynamical behavior which determines the cell fate at different cellular processes levels including cell cycle progression, cell survival, cell death, metabolic reprogramming, and aging. The dynamics of the complex responses to rapamycin in cancer cells have been attributed to its differential time-dependent inhibitory effects on mTORC1 and mTORC2, the two main complexes of mTOR. Two explanations were previously provided for this phenomenon: 1-Rapamycin does not inhibit mTORC2 directly, whereas it prevents mTORC2 formation by sequestering free mTOR protein. 2-Components like Phosphatidic Acid further stabilize mTORC2 compared with mTORC1. To understand the mechanism by which rapamycin differentially inhibits the mTOR complexes, we present a mathematical model of rapamycin mode of action based on the first explanation, i.e., *Le Chatelier’*s principle. Translating the interactions among components of mTORC1 and mTORC2 into a mathematical model revealed the dynamics of rapamycin action in different doses and time-intervals of rapamycin treatment. The model shows that rapamycin has stronger effects on mTORC1 compared with mTORC2, simply due to its direct interaction with free mTOR and mTORC1, but not mTORC2, without the need to consider other components that might further stabilize mTORC2. Based on our results, even when mTORC2 is less stable compared with mTORC1, it can be less inhibited by rapamycin.

## Introduction

Cancer cells have dysregulated networks of signaling pathways which provide them with the capability to develop specific characteristics defined as hallmarks of cancer [1-3]. Considering the functions of these signaling pathways in time and space using mathematical models and systems-level approaches offer novel insights for the discovery and optimization of therapeutic regimens for cancer [4-6]. The ability of cancer cells to develop resistance and escape from effects of treatment modalities comes from the complex information processing system which surfs the stress signals within and among the cancer cells. Systems-level analysis of high-throughput omics data of signal propagation and decision-making system in cancer cells can discover new precision and personalized treatments. Understanding the dynamics of functional modules and signal integrators in malignant cells is an essential part of these efforts [3, 7, 8].

Mammalian target of rapamycin (mTOR) also known as mechanistic target of rapamycin is an important signal integrator in the cells which controls many functions such as cell survival, cell death and aging [9-11]. mTOR is a serine-threonine protein kinase which functions at the intersection of the networks of cellular homeostasis control. mTOR receives signals from different functional modules in cells such as stress sensors, metabolic imbalance sensors, cell cycle, and cell size control system and within its complex interactions with other compartments contribute to cell fate decision in terms of cell death and survival [1, 10-13]. Role of mTOR as a signal integrator not only is defined by the signaling networks in which it is involved but also by the temporal processes of signal integration and transduction [14-17].

Organization of the mTOR signaling pathway has a modular design in which two main complexes, which are mTOR complex 1 (mTORC1) and mTOR Complex 2 (mTORC2), are the basic elements. In terms of responses of mTOR to different signals, mTORC1 controls activation of synthesis of macromolecules such as proteins and lipids, cell growth and cell cycle progression, metabolic re-programming and suppression of autophagy. Basically, under physiological condition, when there is no cellular stress, and cell metabolic state is at homeostasis in the presence of growth factors, sufficient nutrients (particularly amino acids), and enough oxygen, mTORC1 transduces a signal to increase production of cellular building blocks, as well as the cell growth, while suppressing autophagy which degrades the cell’s own materials and organelles. Inhibition of mTOR, which happens under stress condition and drug treatment, induces opposite responses to keep the cell survival and avoid cell death in time of limited nutrients and increased stress [1, 10, 11, 18, 19]. (**Figure 1**)

**Figure 1:**
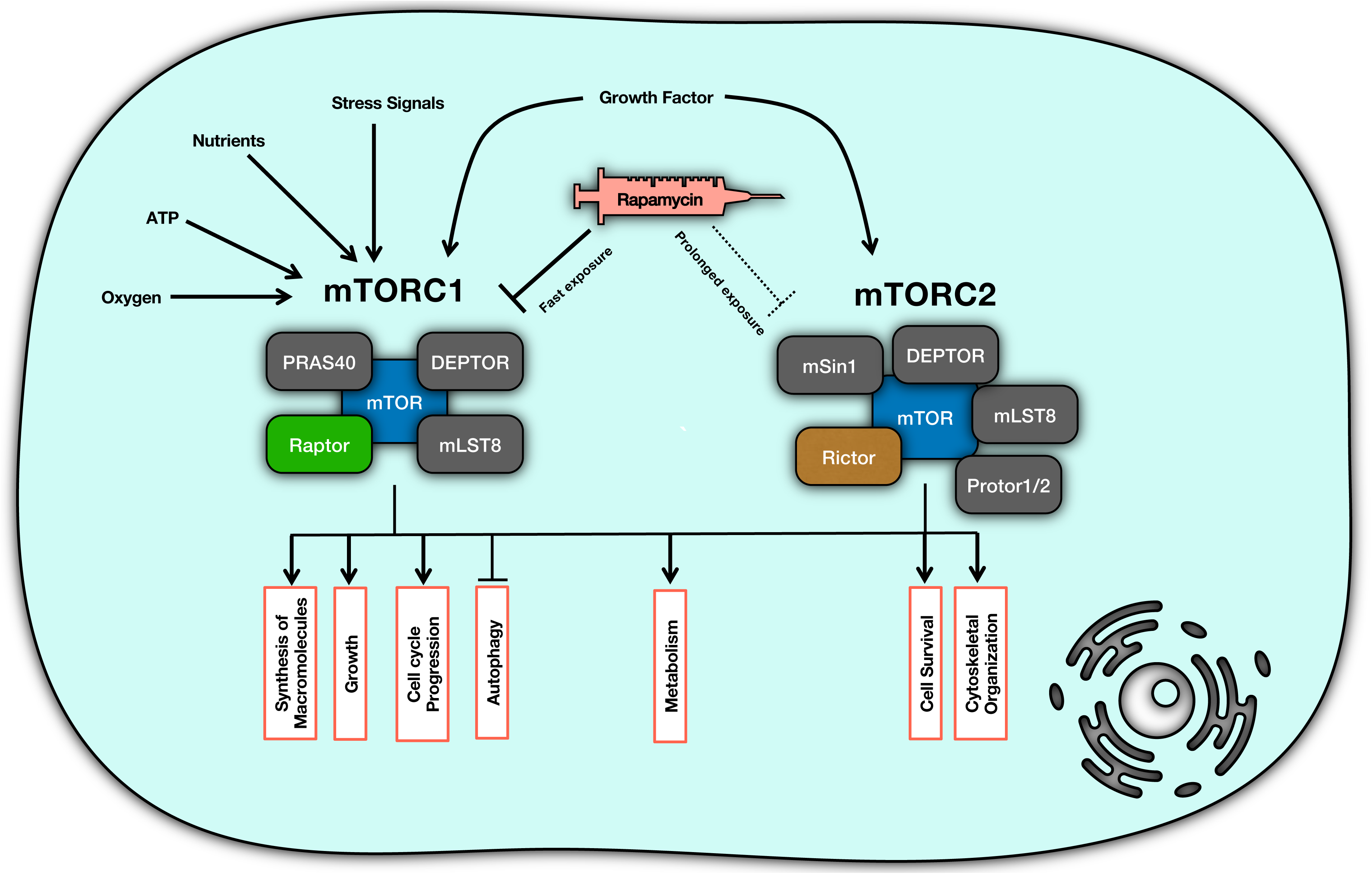
mTOR complex organization and functions. mTOR plays its role in signal integration through two main components of mTORC1 and mTORC2. Each of these components is responsible for transducing appropriate signals into distinct responses.

mTOC1 is a protein complex composed of mTOR, DEPTOR, PRAS40, mLST8, and Raptor [20, 21]; while mTORC2 consists of another set of proteins including mTOR, mSin, DEPTOR, Rictor, Protor1/2 and mLST8 [20, 22]. mTORC2 contributes to metabolic reprogramming, cytoskeletal organization and cell survival through signal transduction when it receives the signal from growth factors [10, 11, 20]. To clarify, mTORC2 phosphorylates Akt (Protein Kinase B) and activates the survival responses downstream of Akt [10, 11, 14, 18].

Rapamycin (sirolimus) is a macrolide with immunosuppressive, anti-neoplastic and anti-aging effects which is known as an mTOR inhibitor [20, 23-25]. Rapamycin is used as an mTOR inhibitor for different clinical conditions such as cancer and organ transplantation [26, 27]. Upon entering a cell, rapamycin binds to 12-kDa FK506-binding protein (FKBP12), and then this complex suppresses the kinase activity of mTORC1 [24]. Although it is believed that rapamycin imposes its inhibition on mTOR via mTORC1 complex [11, 28], but there is evidence showing a more complex mechanism of action which is determined based on the temporal properties of rapamycin treatment [24, 29-31]. For example, short-term treatment with rapamycin renders cells more sensitive to insulin via mTORC1 while long-term exposure increases the insulin resistance which happens via affecting mTORC2 [31]. In fact, prolonged rapamycin treatment can inhibit mTORC2, leading to inhibition of Akt/PKB signal [24]. The topology of molecular interactions in the mTOR pathway cannot provide enough insights on how differential responses can occur with different temporal dose schedules of rapamycin treatment. The dynamics of this interaction network can shed light on paradoxical responses in terms of mTORC1 and mTORC2 activities [32, 33]. To address this question, we used a dynamical model of interactions of the mTOR complexes. There are some facts on time-scales of mTOC1 and mTOC2 behaviors: inhibition of mTORC1 happens in the first 30 minutes of exposure while HeLa cells, for instance, lose the mTORC2 only if the rapamycin exposure takes up to at least 24 hours [24].

We hypothesize that it is based on *Le Chatelier*’s principle [34] that rapamycin can inhibit mTORC2. Rapamycin binds directly to mTORC1 and dissociates Raptor from mTOR, while it is unable to bind to pre-formed mTORC2. Sequestration of mTOR by rapamycin eventually blocks the formation of new mTORC2, and that is why the inhibition of mTORC2 takes much time to happen. (**Figure 2**)

**Figure 2:**
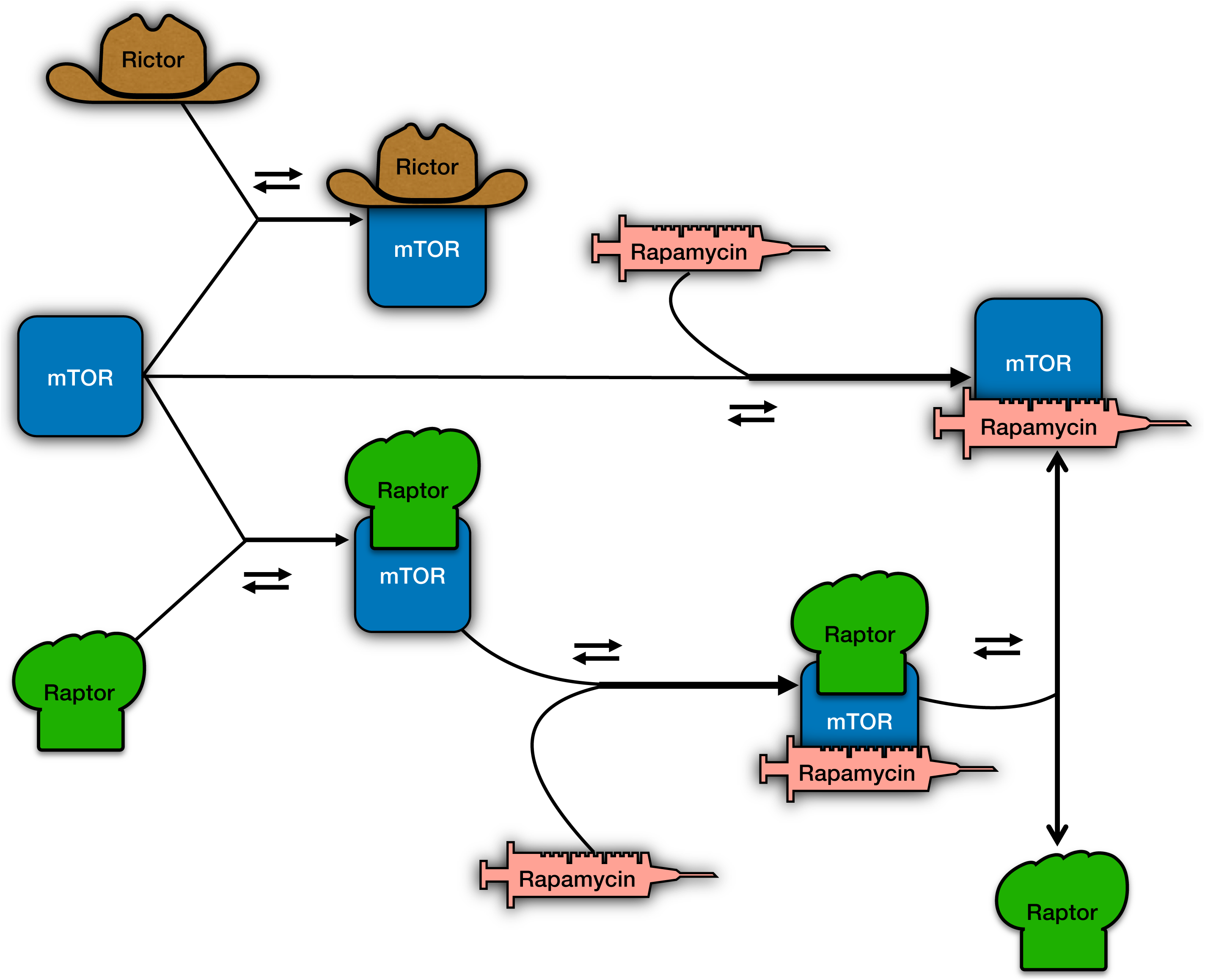
The schematic interaction diagram representing the mechanism of inhibition of mTOR by rapamycin.

Another possible mechanism can be the contribution of Phosphatidic Acid (PA) which is required for the formation of mTORC1 and mTORC2. PA stabilizes mTORC2 more strongly compared to mTORC1 while rapamycin reduces this stability [35, 36]. To analyze the dynamics of rapamycin action upon mTOR complexes and evaluate the validity of this hypothesis, we present a mathematical model of interactions among rapamycin, mTOR, mTORC1, and mTORC2using using Ordinary Differential Equations (ODEs).

## Methods

### The Model

A model of cellular responses to rapamycin using mTOR signaling was developed by translating the interaction diagram in **Figure 2** into a set of ODEs. Since all the reactions in this model are concentration-dependent, they follow the law of mass action and the first order kinetic law. We also assume that we are treating a population of cells that all of them are resistant to rapamycin-induced apoptosis; thus, no cell death occurs after treatment. When administered for a long time, we then assume that when cells divide, all daughter cells carry the same amount of species as the mother cells. Therefore, simulation for one cell represents the result of the average of the cell population, regardless of the number of cell divisions occurred. The list of species and parameters and the final values assigned to them are provided in **Tables 1, 2 and 3**. The model was simulated in the SimBiology application of MATLAB (2017a). The SimBiology file is provided as **Supplementary File 1**.

**Table 1:**
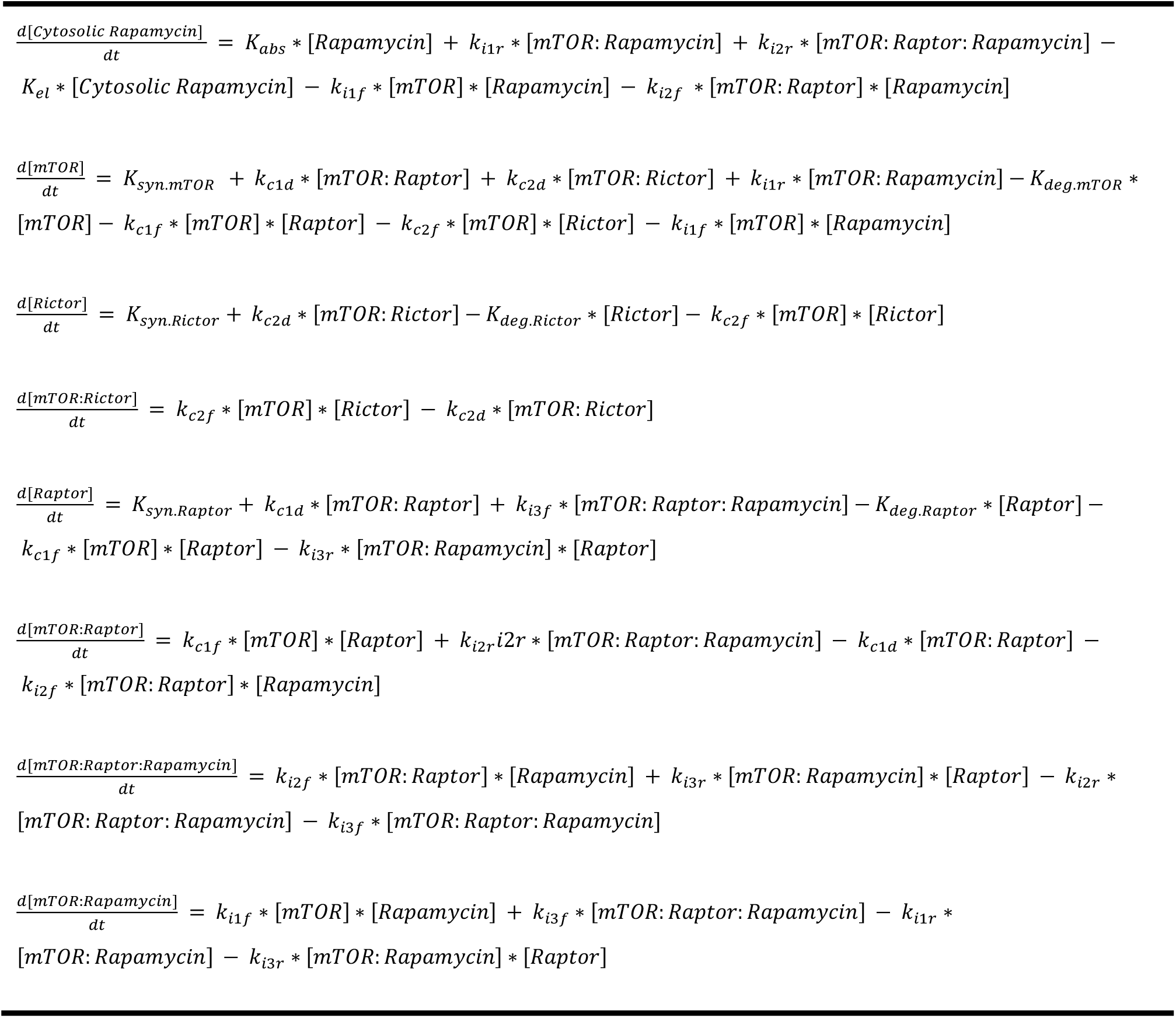
Model Equations

**Table 2:**
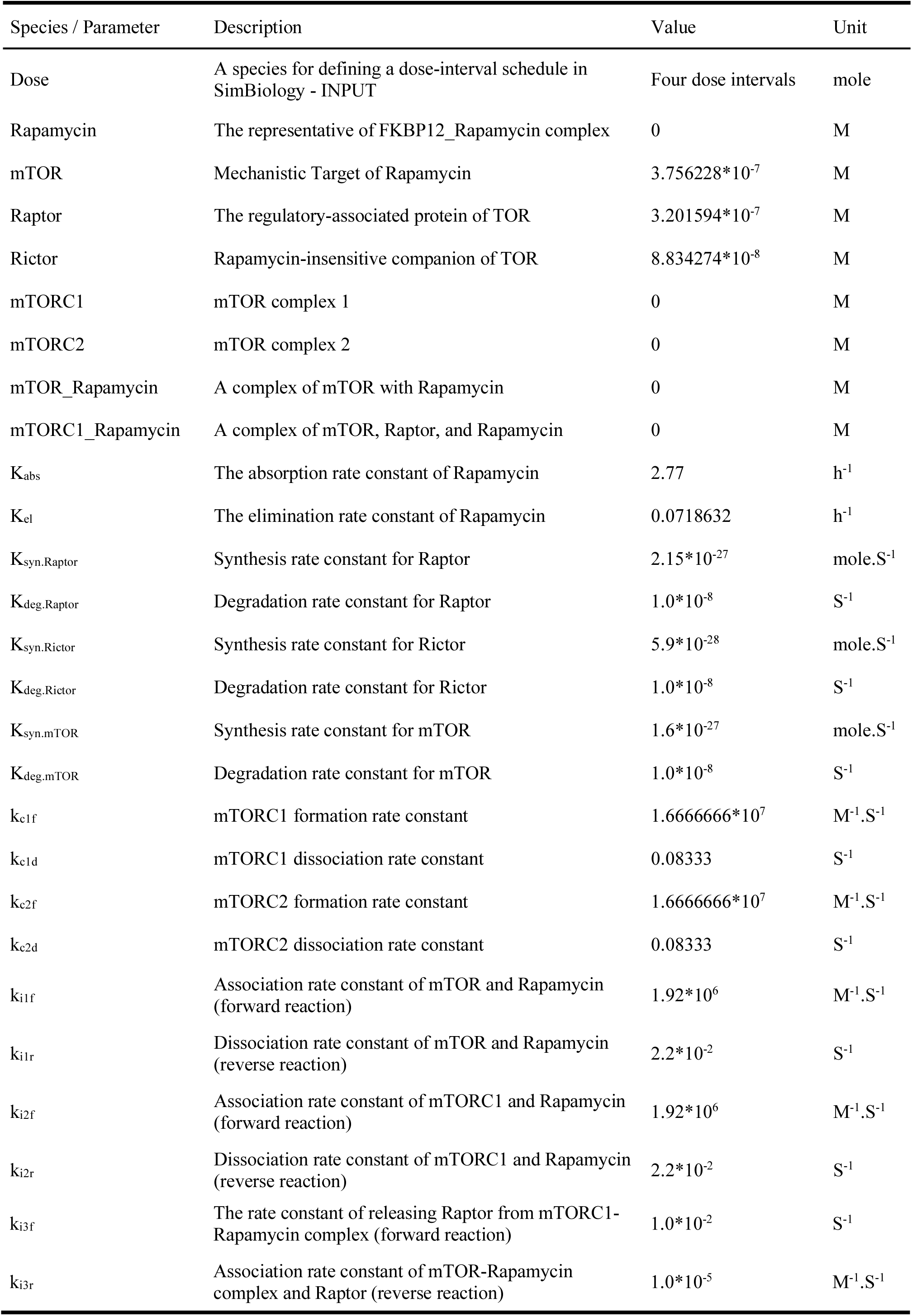
Species and Parameter specifications, their descriptions and initial values used in the model. M= Molar; h^−1^= per hour; mole.S^−1^= mole per Second; M^−1^.S^−1^= per Molar per Second; S^−1^= per Second.

| Species / Parameter | Description | Value | Unit |
| --- | --- | --- | --- |
| Dose | A species for defining a dose-interval schedule in SimBiology - INPUT | Four dose intervals | mole |
| Rapamycin | The representative of FKBP12_Rapamycin complex | 0 | M |
| mTOR | Mechanistic Target of Rapamycin | $3.756228 \times 10^{-7}$ | M |
| Raptor | The regulatory-associated protein of TOR | $3.201594 \times 10^{-7}$ | M |
| Rictor | Rapamycin-insensitive companion of TOR | $8.834274 \times 10^{-8}$ | M |
| mTORC1 | mTOR complex 1 | 0 | M |
| mTORC2 | mTOR complex 2 | 0 | M |
| mTOR_Rapamycin | A complex of mTOR with Rapamycin | 0 | M |
| mTORC1_Rapamycin | A complex of mTOR, Raptor, and Rapamycin | 0 | M |
| K<sub>abs</sub> | The absorption rate constant of Rapamycin | 2.77 | h<sup>-1</sup> |
| K<sub>el</sub> | The elimination rate constant of Rapamycin | 0.0718632 | h<sup>-1</sup> |
| K<sub>syn.Raptor</sub> | Synthesis rate constant for Raptor | $2.15 \times 10^{-27}$ | mole.S<sup>-1</sup> |
| K<sub>deg.Raptor</sub> | Degradation rate constant for Raptor | $1.0 \times 10^{-8}$ | S<sup>-1</sup> |
| K<sub>syn.Rictor</sub> | Synthesis rate constant for Rictor | $5.9 \times 10^{-28}$ | mole.S<sup>-1</sup> |
| K<sub>deg.Rictor</sub> | Degradation rate constant for Rictor | $1.0 \times 10^{-8}$ | S<sup>-1</sup> |
| K<sub>syn.mTOR</sub> | Synthesis rate constant for mTOR | $1.6 \times 10^{-27}$ | mole.S<sup>-1</sup> |
| K<sub>deg.mTOR</sub> | Degradation rate constant for mTOR | $1.0 \times 10^{-8}$ | S<sup>-1</sup> |
| k<sub>c1f</sub> | mTORC1 formation rate constant | $1.6666666 \times 10^7$ | M<sup>-1</sup>.S<sup>-1</sup> |
| k<sub>c1d</sub> | mTORC1 dissociation rate constant | 0.08333 | S<sup>-1</sup> |
| k<sub>c2f</sub> | mTORC2 formation rate constant | $1.6666666 \times 10^7$ | M<sup>-1</sup>.S<sup>-1</sup> |
| k<sub>c2d</sub> | mTORC2 dissociation rate constant | 0.08333 | S<sup>-1</sup> |
| k<sub>i1f</sub> | Association rate constant of mTOR and Rapamycin (forward reaction) | $1.92 \times 10^6$ | M<sup>-1</sup>.S<sup>-1</sup> |
| k<sub>i1r</sub> | Dissociation rate constant of mTOR and Rapamycin (reverse reaction) | $2.2 \times 10^{-2}$ | S<sup>-1</sup> |
| k<sub>i2f</sub> | Association rate constant of mTORC1 and Rapamycin (forward reaction) | $1.92 \times 10^6$ | M<sup>-1</sup>.S<sup>-1</sup> |
| k<sub>i2r</sub> | Dissociation rate constant of mTORC1 and Rapamycin (reverse reaction) | $2.2 \times 10^{-2}$ | S<sup>-1</sup> |
| $k_{i3f}$ | The rate constant of releasing Raptor from mTORC1-Rapamycin complex (forward reaction) | $1.0 \cdot 10^{-2}$ | $S^{-1}$ |
| $k_{i3r}$ | Association rate constant of mTOR-Rapamycin complex and Raptor (reverse reaction) | $1.0 \cdot 10^{-5}$ | $M^{-1} \cdot S^{-1}$ |

**Table 3:** List of reactions of the model with their rate constants and their reaction rate equation.

| Reaction | Rate Constant | Reaction rate | Description |
| --- | --- | --- | --- |
| Dose → Rapamycin | $K_{abs}$ | $K_{abs} \cdot \text{Dose}$ | Absorption of rapamycin |
| Rapamycin → null | $K_{el}$ | $K_{el} \cdot [\text{Rapamycin}]$ | Elimination of rapamycin |
| null → mTOR | $K_{syn.mTOR}$ | $K_{syn.mTOR}$ | Synthesis of mTOR |
| mTOR → | $K_{deg.mTOR}$ | $K_{deg.mTOR} \cdot [\text{mTOR}]$ | Degradation of mTOR |
| → Raptor | $K_{syn.Raptor}$ | $K_{syn.Raptor}$ | Synthesis of Raptor |
| Raptor → | $K_{deg.Raptor}$ | $K_{deg.Raptor} \cdot [\text{Raptor}]$ | Degradation of Raptor |
| → Rictor | $K_{syn.Rictor}$ | $K_{syn.Rictor}$ | Synthesis of Rictor |
| Rictor → | $K_{deg.Rictor}$ | $K_{deg.Rictor} \cdot [\text{Rictor}]$ | Degradation of Rictor |
| mTOR + Raptor ⇌ mTORC1 | $k_{c1f}, k_{c1d}$ | $k_{c1f} \cdot [\text{mTOR}] \cdot [\text{Raptor}] - k_{c1d} \cdot [\text{mTORC1}]$ | mTORC1 formation and dissociation |
| mTOR + Rictor ⇌ mTORC2 | $k_{c2f}, k_{c2d}$ | $k_{c2f} \cdot [\text{mTOR}] \cdot [\text{Rictor}] - k_{c2d} \cdot [\text{mTORC2}]$ | mTORC2 formation and dissociation |
| mTOR + Rapamycin ⇌ mTOR_Rapamycin | $k_{i1f}, k_{i1r}$ | $k_{i1f} \cdot [\text{mTOR}] \cdot [\text{Rapamycin}] - k_{i1r} \cdot [\text{mTOR\_Rapamycin}]$ | mTOR - Rapamycin association/sequestration and dissociation |
| mTORC1 + Rapamycin ⇌ mTORC1_Rapamycin | $k_{i2f}, k_{i2r}$ | $k_{i2f} \cdot [\text{mTORC1}] \cdot [\text{Rapamycin}] - k_{i2r} \cdot [\text{mTORC1\_Rapamycin}]$ | mTORC1 - Rapamycin association/sequestration and dissociation |
| mTORC1_Rapamycin ⇌ mTOR_Rapamycin + Raptor | $k_{i3f}, k_{i3r}$ | $k_{i3f} \cdot [\text{mTORC1\_Rapamycin}] - k_{i3r} \cdot [\text{mTOR\_Rapamycin}] \cdot [\text{Raptor}]$ | Releasing Raptor from mTORC1 - rapamycin complex |

### Parameters

We extracted kinetic parameters of rapamycin (K_abs_ = 2.77 h^−1^, A = 7.89 ng/ml, B = 1.06 ng/ml, **α** = 0.28 h^−1^ and **β** = 0.011 h^−1^) from Ferron et al. [37]. They are listed in **Supplementary Table S1**. K_el_ = 0.0718632 h^−1^ was calculated as: 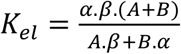 [37]. Moreover, association and dissociation rate constants of mTOR and Raptor were extracted from Vinod and Venkatesh [38]. They assumed the association rate constant of Raptor and mTOR to be 1 nM^−1^.min^−1^ (= 1.6*10^7^ M^−1^.S^−1^) and calculated the dissociation rate constant to be 5 min^−1^ (= 0.083 S^−1^) [38]. (**Supplementary Table S2**)

Association and dissociation rate constants of rapamycin and mTOR were extracted from Banaszynski et al. [39], which reported the association and dissociation rate constants of formation of trimeric FKBP12-rapamycin-mTOR complex as 1.92*10^6^ M^−1^.S^−1^ and 2.2*10^−2^ S^−1^, respectively. Hereby, for the simplicity of our model, we ignored the binding of rapamycin with FKBP12 and assumed the cytosolic rapamycin is FKBP12-Rapamycin (**Supplementary Table S2**).

Similarly, the average quantity of mTOR, Raptor, and Rictor in a cell was extracted from Geiger et al. [40] which measured total proteome of eleven cell lines. Furthermore, we assumed the volume of a cell to be 5*10^−12^ liter and calculated concentrations of mTOR, Raptor, and Rictor based on Geiger et al.[40]. (**Supplementary Table S2**)

We could not find any studies that even could remotely estimate mTOR-Rictor binding rate constants. Hence, initially, we assumed the association-dissociation rate constants of mTOR and Rictor interaction (k_C2f_ and k_C2d_) to be equal to that of mTOR and Raptor interaction (k_C1f_ and k_C1d_). Then we scanned every parameter space within a range logarithmically spaced from −3logs to +3logs using seven steps to see how changing each parameter affects the outputs. Additionally, we scanned the parameter space of the macro-constants of rapamycin (K_abs_ and K_el_) within a range logarithmically spaced from −2logs to +2logs using three steps.

Moreover, we chose the values for synthesis and degradation rate constants of mTOR, Raptor, and Rictor in a way to have the least effect on the main question, also, in a way to have an almost similar amount of these species, in total, in the beginning, and after reaching the equilibrium.

### Dose schedules for rapamycin

Four dose-interval schedules were selected to be used in this model:

I. A low-dose and short-interval schedule (8.0*10^−20^ mole/Day).
II. A high-dose and short-interval schedule (5.6*10^−19^ mole/Day).
III. A high-dose and long-interval schedule (5.6*10^−19^ mole/Week).
IV. A very high-dose and long-interval schedule (2.24*10^−18^ mole/Week).

Considering the maximum whole-blood concentration of rapamycin for transplant patients equal to ∼15 ng/ml [41] and the molecular weight of rapamycin equal to ∼914 g/mole [42], if we assume the cytosolic concentration of rapamycin is equal to that of the whole-blood, we can calculate that ∼8.0 × 10^−20^ moles of rapamycin is present in a cell with the volume of 5.0 × 10^−12^ liter. We implemented this amount as the lowest daily dose (regimen I). By the same method, if we want to have a weekly schedule with the same cumulative dose as regimen I, we have 5.6 × 10^−19^ mole/Week (regimen III). Moreover, we can administer 5.6 × 10^−19^ moles of rapamycin on a daily basis to have regimen II. Finally, we can multiply the third regimen by 4 to have 2.24 × 10^−18^ mole/Week as a very high dose with a weekly interval (regimen IV). Besides this, we scanned a ± 20% area (linearly divided into five steps) for every treatment we defined in the simulations. The dose specifications in SimBiology are listed in **Table 4**.

**Table 4:** Dose-interval specifications

| Description | Name | Start Time | Amount | Interval | Repeat Count | Area Scanned ($\pm 20\%$) |
| --- | --- | --- | --- | --- | --- | --- |
| Low-dose and short-interval | $8.0 \times 10^{-20}$ mole/Day | 240 h | $8.0 \times 10^{-20}$ moles | 24 h | 95 | $6.4 \times 10^{-20}$ to $9.6 \times 10^{-20}$ moles |
| High-dose and short-interval | $5.6 \times 10^{-19}$ mole/Day | 240 h | $5.6 \times 10^{-19}$ moles | 24 h | 95 | $4.48 \times 10^{-19}$ to $6.72 \times 10^{-19}$ moles |
| High-dose and long-interval | $5.6 \times 10^{-19}$ mole/Week | 240 h | $5.6 \times 10^{-19}$ moles | 168 h | 15 | $4.48 \times 10^{-19}$ to $6.72 \times 10^{-19}$ moles |
| Very High-dose and long-interval | $2.24 \times 10^{-18}$ mole/Week | 240 h | $2.24 \times 10^{-18}$ moles | 168 h | 15 | $1.792 \times 10^{-18}$ to $2.688 \times 10^{-18}$ moles |

We measured minimum, maximum and average residual active percentage of mTORC1 (mTORC1%_min_, mTORC1%_max,_ and mTORC1%_avg._) and mTORC2 (mTORC2%_min_, mTORC2%_max,_ and mTORC2%_avg._) in each schedule as a representative of their activity.

## Results

The mathematical model based on the interaction diagram of **Figure 2**, can generate temporal changes in the concentration of different components in the model and reveal the dynamics of the mode of action of rapamycin using different doses of rapamycin within different treatment schedules. The temporal changes of rapamycin concentration in regimens I to IV are shown in **Figure 3**. The corresponding temporal concentration of mTORC1 and mTORC2 is presented in **Figure 4**, and the corresponding data can be found in **Table 5** as the minimum, average and maximum residual active percentage of the two complexes, for the exact 100% dose of each schedule. In **Supplementary Table S3**, the minimum, and maximum of the residual active concentration and remaining active percentage of the two complexes, corresponding to the scanned dose of each schedule are presented. With implementing the initial values of parameters into the model, minimum, maximum and average residual active percentage of mTORC2 is always higher than mTORC1 for all four dose-interval schedules (**Table 5**). Comparing the regimen I and III, which have equal cumulative doses, we observe that rapamycin inhibits mTORC1 to an almost identical extent on average (mTORC1%_avg._ in regimen I and III are 59.26% and 58.01%, respectively). The difference is that the percentage of active mTORC1 in regimen III has a broader range, compared to mTORC1 in regimen I (57.32% to 61.20% vs. 43.72% to 72.30%). A similar phenomenon happens regarding mTORC2 as well (comparing regimen I and III): an average of 83.53% of mTORC2 is remaining active in regimen I with a range of 82.51% to 84.54%, versus an average of 82.02% with a range of 74.33% to 89.71% in regimen III (**Table 5**).

**Table 5:** Comparing the remaining active percent of mTORC1 and mTORC2 based on exact (100%) dose of each schedule. The minimum and maximum values are rounded to two decimal points, then the average is calculated.

|  |  | mTORC1% | mTORC2% |
| --- | --- | --- | --- |
| Drug-free period |  | 100% | 100% |
| 8.0*10<sup>-20</sup> mole/Day | min | 57.32 | 82.51 |
|  | Avg. | 59.26 | 83.53 |
|  | max | 61.20 | 84.54 |
| 5.6*10<sup>-19</sup> mole/Day | min | 12.18 | 43.95 |
|  | Avg. | 18.28 | 51.03 |
|  | max | 24.37 | 58.12 |
| 5.6*10<sup>-19</sup> mole/Week | min | 43.72 | 74.33 |
|  | Avg. | 58.01 | 82.02 |
|  | max | 72.30 | 89.71 |
| 2.24*10<sup>-18</sup> mole/Week | min | 5.43 | 32.72 |
|  | Avg. | 33.37 | 58.43 |
|  | max | 61.30 | 84.14 |

**Figure 3:**
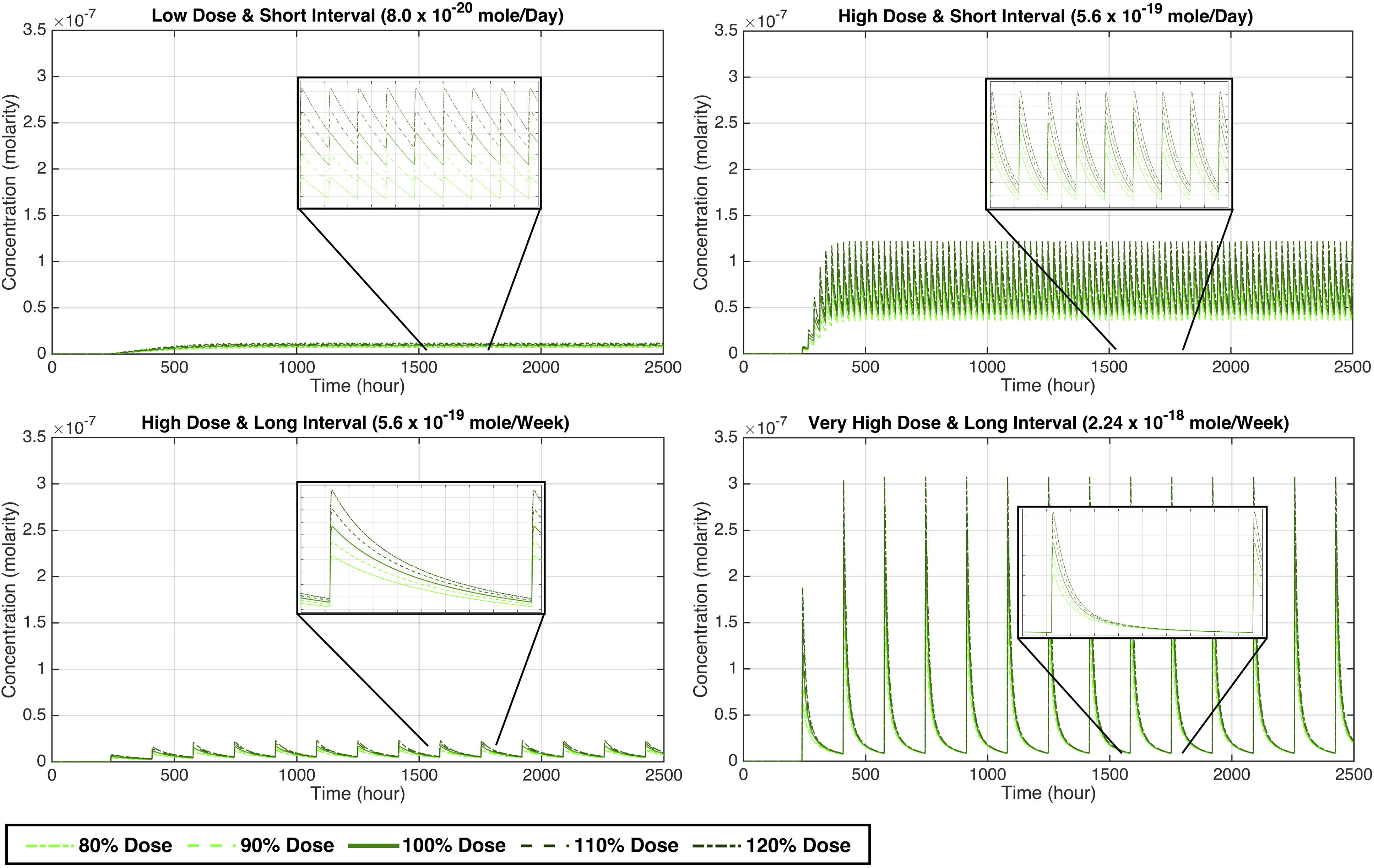
Temporal concentration of rapamycin for regimens I to IV. Each regimen is simulated while scanning the “amount ± 20%” area (linearly stepped) around the dose administered. A) Regimen I: 8.0 * 10^−20^ ± 20% mole/Day; B) Regimen II: 5.6 * 10^−19^ ± 20% mole/Day; C) Regimen III: 5.6 * 10^−19^ ± 20% mole/Week; and D) Regimen IV: 2.24 * 10^−18^ ± 20% mole/ Week.

**Figure 4:**
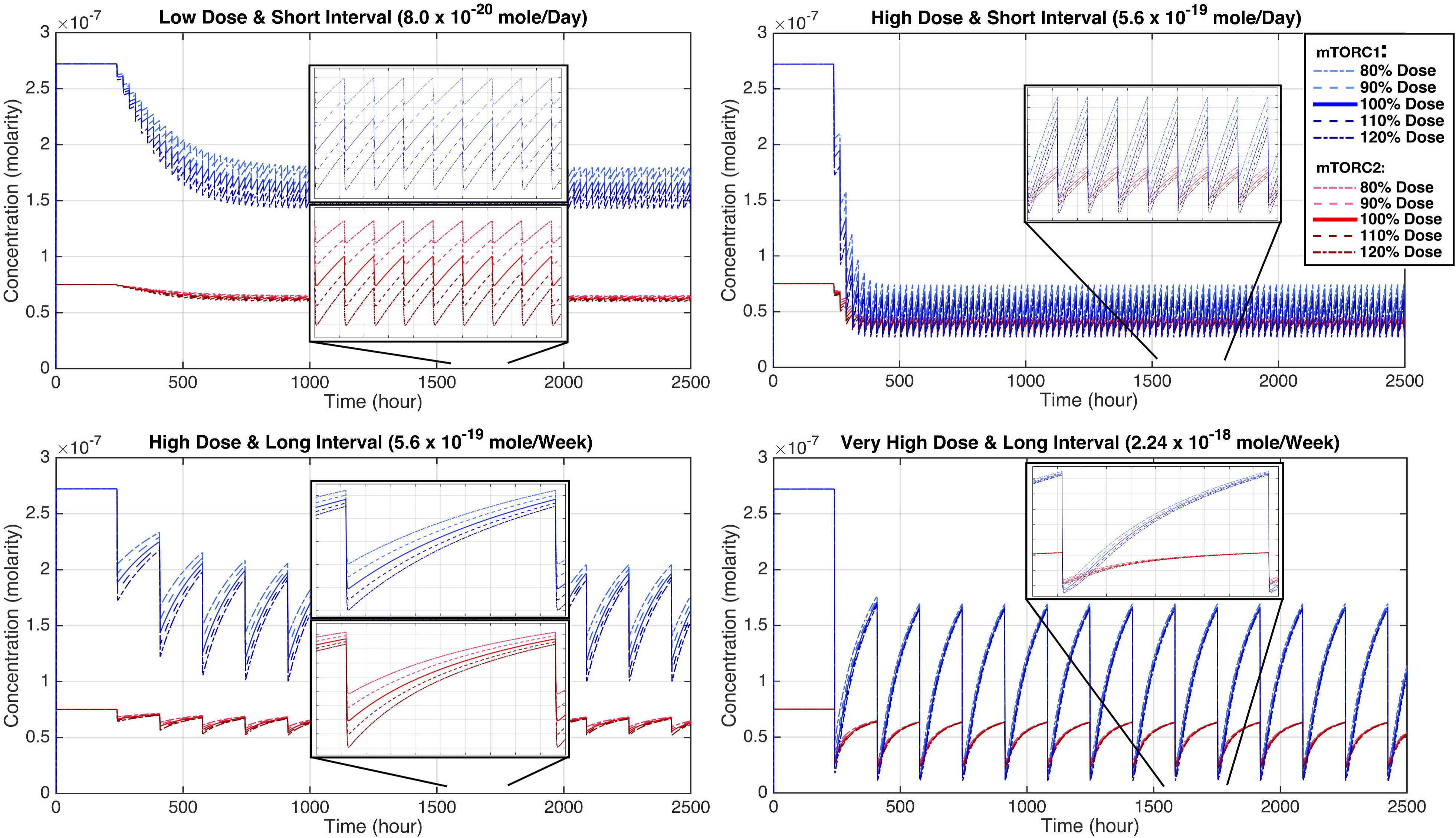
Temporal concentration of mTORC1 and mTORC2 for regimens I to IV. These plots display changes of [mTORC1] and [mTORC2], simultaneously, during the dose administration. Each regimen is simulated while scanning the “amount ± 20%” area (linearly stepped) around the dose administered (the corresponding dose is shown in **Figure 3**). A) Regimen I: 8.0 × 10^−20^ ± 20% mole/Day; B) Regimen II: 5.6 × 10^−19^ ± 20% mole/Day; C) Regimen III: 5.6 × 10^−19^ ± 20% mole/Week; and D) Regimen IV: 2.24 × 10^−18^ ± 20% mole/ Week.

Furthermore, the distinguishable feature of the regimen II is that not only it has a very high C_max_ concentration but also possesses a much higher pre-dose concentration of rapamycin, in contrast to every other schedule (**Figure 3**). Also, both mTOR complexes are almost strongly inhibited by rapamycin in this schedule (regimen II): mTORC1%_avg._ equals 18.28% and mTORC2%_avg._ is 51.03% (**Figure 4** and **Table 5**). In an almost similar fashion, the regimen IV has a very high C_max_ and a high enough C_min_ (higher than C_min_ of the regimen I and III, but not regimen II) that keeps mTORC1 inhibited, even in the pre-dose time (61.30% residual activity) while mTORC2 activity comes back to almost normal (84.14% residual activity).

Altering both K_abs_ and K_el_ in two orders of magnitude (± 2logs) changes the temporal concentration of rapamycin, mTORC1, and mTORC2. However, mTORC1 is still inhibited much more profoundly compared with mTORC2. (**Figure 5 and Supplementary Table S4**)

**Figure 5:**
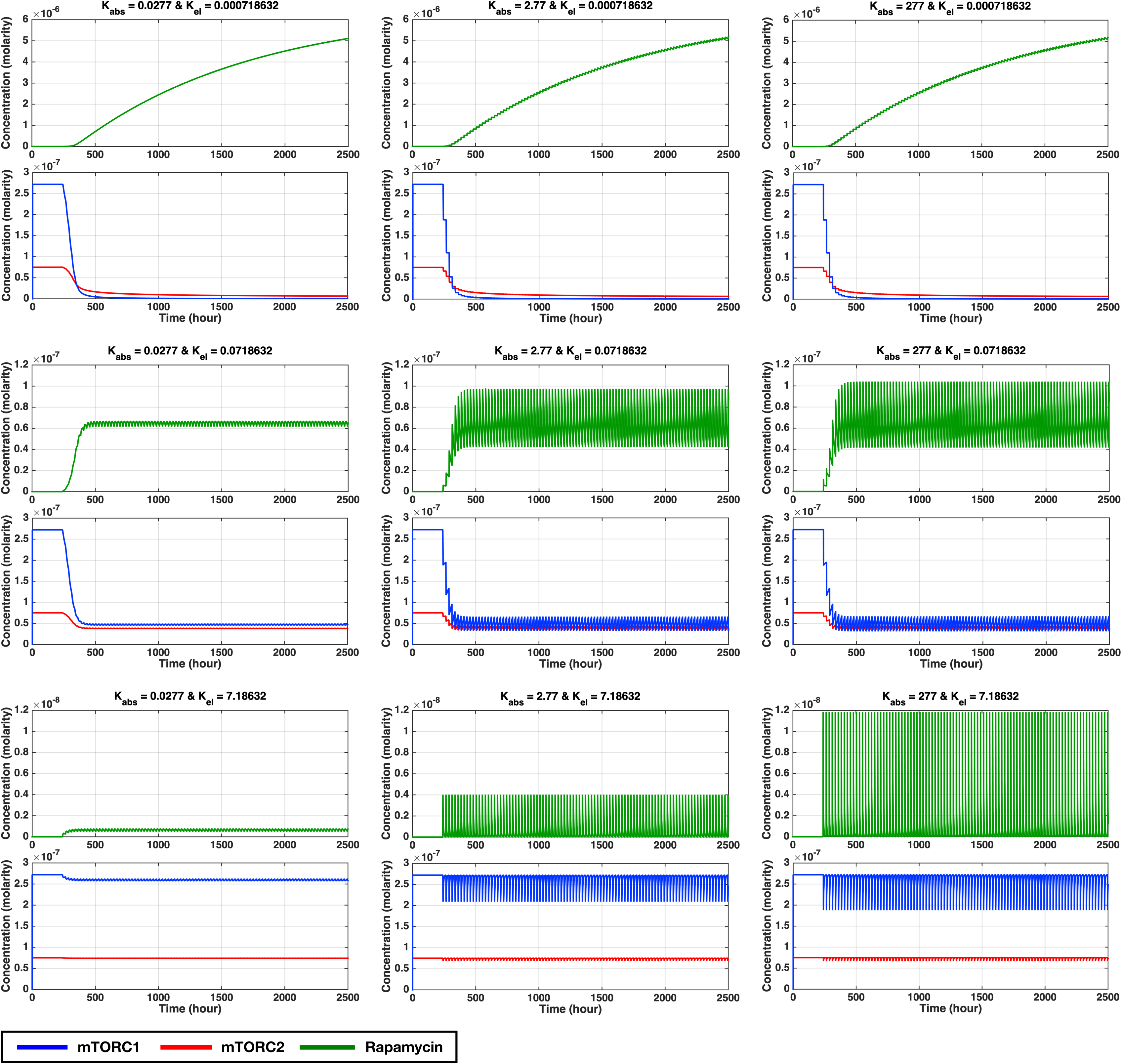
Temporal concentration of rapamycin, mTORC1, and mTORC2 for different absorption and elimination rate constants (macro-constants) of rapamycin: the K_abs_ and K_el_ were scanned jointly in an area within a percentage range logarithmically spaced from −2 to +2 using three steps each, making nine possible combinations for these kinetic parameters. The reference dose used for this parameter scan is 5.0*10^−19^ mole/Day (regimen II). The plot at the center is where the K_abs_ and K_el_ are at their initial values. (k_abs_ = absorption rate constant; K_el_ = elimination rate constant)

The mTORC1%_avg._ and mTORC2%_avg._ are affected by changing the value of formation and dissociation rate constants of these two complexes (k_c1f_, k_c1d_, k_c2f,_ and k_c2d_) in an area of ± 3logs (**Figure S1 and Table 6**). Note that if the C2% > C1% but K_D1_< K_D2_, it means that while mTORC2 is weaker than mTORC1, mTORC2 is less affected by rapamycin.

**Table 6:** Average percent of remaining active mTORC1 and mTORC2 in the parameter scan for association and dissociation rate constants (k_c1f_, k_c1d_, k_c2f_, and k_c2d)_. Every rate constant was scanned separately in an area of ± 3logs in 7 steps. The reference dose is 5.6*10^−19^ mole/day (regimen II). k_c1f_ is the mTORC1 formation rate constant. k_c1d_ is the mTORC1 dissociation rate constant. k_c2f_ is the mTORC2 formation rate constant. k_c2d_ is the mTORC2 dissociation rate constant. C1%_avg._ is the average residual active percentage of mTORC1. C2%_avg._ is the average residual active percentage of mTORC2. K_D1_ is the equilibrium dissociation constant of mTORC1. K_D2_ is the equilibrium dissociation constant of mTORC2.

| | | $C1\%_{avg.}$ | $C2\%_{avg.}$ | $K_{D1}$ | $K_{D2}$ | |
| --- | --- | --- | --- | --- | --- | --- |
| $k_{c1f} = 1.6666666 \cdot 10^7$ | -3Logs | 11.40 | 90.90 | $4.9999 \cdot 10^{-6}$ | $4.9999 \cdot 10^{-9}$ | |
| | -2Logs | 13.62 | 88.84 | $4.9999 \cdot 10^{-7}$ | $4.9999 \cdot 10^{-9}$ | |
| | -Log | 16.68 | 78.38 | $4.9999 \cdot 10^{-8}$ | $4.9999 \cdot 10^{-9}$ | |
| | 1 | 18.27 | 51.04 | $4.9999 \cdot 10^{-9}$ | $4.9999 \cdot 10^{-9}$ | While $K_{D1}$ is not greater than $K_{D2}$, mTORC2 is inhibited less than mTORC1. |
| | Log | 18.83 | 19.07 | $4.9999 \cdot 10^{-10}$ | $4.9999 \cdot 10^{-9}$ | While $K_{D1}$ is not greater than $K_{D2}$, mTORC2 is inhibited less than mTORC1. |
| | 2Logs | 18.91 | 3.33 | $4.9999 \cdot 10^{-11}$ | $4.9999 \cdot 10^{-9}$ | |
| | 3Logs | 18.92 | 0.37 | $4.9999 \cdot 10^{-12}$ | $4.9999 \cdot 10^{-9}$ | |
| $k_{c1d} = 0.08333$ | -3Logs | 17.25 | 46.54 | $4.9999 \cdot 10^{-12}$ | $4.9999 \cdot 10^{-9}$ | While $K_{D1}$ is not greater than $K_{D2}$, mTORC2 is inhibited less than mTORC1 |
| | -2Logs | 17.33 | 44.69 | $4.9999 \cdot 10^{-11}$ | $4.9999 \cdot 10^{-9}$ | While $K_{D1}$ is not greater than $K_{D2}$, mTORC2 is inhibited less than mTORC1 |
| | -Log | 17.77 | 42.10 | $4.9999 \cdot 10^{-10}$ | $4.9999 \cdot 10^{-9}$ | While $K_{D1}$ is not greater than $K_{D2}$, mTORC2 is inhibited less than mTORC1 |
| | 1 | 18.28 | 51.03 | $4.9999 \cdot 10^{-9}$ | $4.9999 \cdot 10^{-9}$ | While $K_{D1}$ is not greater than $K_{D2}$, mTORC2 is inhibited less than mTORC1 |
| | Log | 17.99 | 74.58 | $4.9999 \cdot 10^{-8}$ | $4.9999 \cdot 10^{-9}$ | |
| | 2Logs | 16.85 | 87.58 | $4.9999 \cdot 10^{-7}$ | $4.9999 \cdot 10^{-9}$ | |
| | 3Logs | 15.69 | 90.67 | $4.9999 \cdot 10^{-6}$ | $4.9999 \cdot 10^{-9}$ | |
| $k_{c2f} = 1.6666666 \cdot 10^7$ | -3Logs | 18.16 | 7.59 | $4.9999 \cdot 10^{-9}$ | $4.9999 \cdot 10^{-6}$ | |
| | -2Logs | 18.24 | 9.19 | $4.9999 \cdot 10^{-9}$ | $4.9999 \cdot 10^{-7}$ | |
| | -Log | 18.51 | 19.43 | $4.9999 \cdot 10^{-9}$ | $4.9999 \cdot 10^{-8}$ | While $K_{D1}$ is not greater than $K_{D2}$, mTORC2 is inhibited less than mTORC1 |
| | 1 | 24.37 | 51.04 | $4.9999 \cdot 10^{-9}$ | $4.9999 \cdot 10^{-9}$ | While $K_{D1}$ is not greater than $K_{D2}$, mTORC2 is inhibited less than mTORC1 |
| | Log | 17.23 | 92.84 | $4.9999 \cdot 10^{-9}$ | $4.9999 \cdot 10^{-10}$ | |
| | 2Logs | 16.85 | 99.27 | $4.9999 \cdot 10^{-9}$ | $4.9999 \cdot 10^{-11}$ | |
| | 3Logs | 16.79 | 99.81 | $4.9999 \cdot 10^{-9}$ | $4.9999 \cdot 10^{-12}$ | |
| $k_{c2d} = 0.08333$ | -3Logs | 16.80 | 99.79 | $4.9999 \cdot 10^{-9}$ | $4.9999 \cdot 10^{-12}$ | |
| | -2Logs | 16.85 | 99.25 | $4.9999 \cdot 10^{-9}$ | $4.9999 \cdot 10^{-11}$ | |
| | -Log | 17.23 | 87.03 | $4.9999 \cdot 10^{-9}$ | $4.9999 \cdot 10^{-10}$ | |
| | 1 | 18.28 | 51.04 | $4.9999 \cdot 10^{-9}$ | $4.9999 \cdot 10^{-9}$ | While $K_{D1}$ is not greater than $K_{D2}$, mTORC2 is inhibited less than mTORC1 |
| | Log | 18.51 | 19.43 | $4.9999 \cdot 10^{-9}$ | $4.9999 \cdot 10^{-8}$ | While $K_{D1}$ is not greater than $K_{D2}$, mTORC2 is inhibited less than mTORC1 |
| | 2Logs | 18.24 | 9.20 | $4.9999 \cdot 10^{-9}$ | $4.9999 \cdot 10^{-7}$ | |
| | 3Logs | 18.16 | 0.76 | $4.9999 \cdot 10^{-9}$ | $4.9999 \cdot 10^{-6}$ | |

**In figure S1 And Supplementary Table S5**, it is shown that how mTORC1%_avg._ and mTORC2%_avg._ are affected by scanning k_i1f_, k_i1r_, k_i2f_, k_i2r_, k_i3f_ and k_i3r_ in an area of ± 3logs. C2% > C1% is true for all cases, which means none of these ± 3logs-scans made mTORC2 being affected more than mTORC1.

## Discussion

The dynamics portrayed by mathematical model shows that even with neglecting the role of PA and only by considering the *Le Chatelier*’s principle [34, 43], mTORC2 is mechanistically inhibited only after prolonged rapamycin exposure in cells. This model magnifies the importance of *in silico* dynamical models to understand the mechanism of signal transduction considering drug treatments. By evaluating the interaction network statically controlling mTOR complexes, rapamycin cannot directly inhibit mTORC2. by observing the dynamics of this network, if the concentration of rapamycin persistently remains high enough, sequestration of the free mTOR prompts the equilibrium of mTORC2 formation (mTOR + Rictor ⇌ mTORC2) to choose the reverse path, based on *Le Chatelier*’s principle, and consumes the pre-formed mTORC2. Based on comprehending our results in conjunction with the literature, PA might only increase the confidence of this mechanism. It means that the amount of PA in a cell only determines how long it takes for the mTORC2 in that specific cell to be affected by rapamycin. According to our mathematical model, mTORC2 inhibition can be avoided by increasing the interval of rapamycin to reduce the trough concentration below the minimum effective dose.

One of the paradoxes regarding the role of mTOR signaling in aging is that rapamycin increases female but not male lifespan in mice [44, 45]. However, Leontieva et al. [46] indicated that “weekly” administration of rapamycin could extend the male lifespan while avoiding the detrimental side effects that other studies described. Regarding metabolic effects of rapamycin, Arriola Apelo et al. [47] clarified that an intermittent administration of rapamycin (every five days) reduces its side effects on glucose metabolism, insulin sensitivity, and immune system. They showed that the reason for this phenomenon is that mTORC2 is not affected in this regimen [47]. They also reported the efficacy of intermittent rapamycin administration in promoting the female lifespan in mice [48]. Again, our model demonstrates the mechanism by which this intermittent administration (long interval) helps in avoiding mTORC2 inhibition.

It has been shown that selective or targeted mTORC2 inhibition causes reduced cell proliferation, tumor size, and angiogenesis in several types of cancers including colon cancer, breast cancer and multiple myeloma [49-53]. Differential effects of dose-interval schedules of rapamycin (or rapalogs) on colon cancer have also been investigated. Guba et al. [54, 55] employed three schedules of rapamycin with equal cumulative doses: I) Once daily (QD); II) Once every three days (Q3D); and III) Continuous IV infusion. Interestingly, they observed that while the IV infusion produced the least plasma concentration, it was the most effective one against colon cancer xenografts. Furthermore, the Q3D dose showed the least efficacy regarding the inhibition of tumor growth. It can be reasonably deduced from this study that prolonged duration of exposure to sufficiently high concentrations of rapamycin is necessary for its mTORC2-related effects. Considering results of Guba et al. [54, 55], comparing regimen I and III of our model (which have equal cumulative doses) clearly shows why the Q3D schedule cannot inhibit tumor growth: if the cumulative dose is constant and the interval is prolonged, while the C_max_ gets higher, the C_min_ (trough concentration) drops drastically, and the mTORC1 and mTORC2 activity comes back to their approximate normal state long before the next dose is administered. Now, not only the mTORC2 signal is not interrupted, but also the negative feedback of mTORC1 on the input signal of growth factors is removed; thus, the tumor cells have the chance to grow and divide again.

It is shown that inhibition of mTORC1 leads to improved immune profiles in the elderly while inhibition of mTORC2 is responsible for adverse events [56]. Mannick et al. [56] have mentioned that mTORC1 inhibition is enough to strengthen the immune system, amplifying the response to influenza vaccine and decreasing the overall rate of infection in elderly subjects [56]. Mannick et al. [57] administered three dose-interval schedules of everolimus, which is a rapamycin-like mTOR inhibitor, (0.5 mg/Day, 5mg/Week, 20mg/Week; like regimens I, III and IV in our model) to elderly subjects to compare their differential effect on responsiveness to influenza vaccine; the 0.5 mg/Day and 5 mg/Week regimens were more successful in amelioration of immunosenescence, while 20 mg/Week regimen showed slightly less efficacy and more side effects. These results could be explained by the lower trough concentrations of everolimus in 0.5 mg/Day and 5 mg/Week schedules, which leads to only partial inhibition of mTORC1 and probably much less inhibition of mTORC2. However, the 20 mg/Week regimen showed more side effects, likely due to almost full inhibition of mTORC1 and (at least) partial inhibition of mTORC2 [57]. Here in our model, comparing regimens III and IV, the activity of both complexes come back to an almost equal amount at the pre-dose area (mTORC1%_max_ = 72.30% vs. 61.30% and mTORC2%_max_ = 89.71% vs. 84.14% for regimen III vs. IV, respectively). However, looking at the average residual activity in these two regimens, both mTOR complexes are much more prominently inhibited in regimen IV (mTORC1%_avg._= 58.01% vs. 33.37% and mTORC2%_avg._= 82.02% vs. 58.43% for regimen III vs. IV, respectively). Hence, our model confirms the findings of Mannick et al. [57]: A very high dose and weekly-administered rapamycin inhibits mTORC2 to a greater extent, leading to extra side effects.

Regarding the association and dissociation rate constants of mTORC2 (k_c2f_ and k_c2d_), and also K_D1_ and K_D2_, it is difficult to conclude which values are the closest to the real values. Toschi et al. [58] claim that K_D1_ must be higher than K_D2_ (meaning, Rictor binds to mTOR more tightly than Raptor does) to see rapamycin inhibits mTORC1 more than mTORC2, and this is made possible by the higher affinity of PA for mTORC2 compared to mTORC1. Simulations of our mathematical model show that even with hypothetical values for mTORC2 rate constants, in some cases, we observe that K_D2_ is less than K_D1_ (which means Raptor has a higher affinity towards mTOR compared to Rictor), yet mTORC1 is still inhibited more than mTORC2. Although we need an accurate experimental design to find out mTORC2 formation and dissociation rate constants, these counterexamples undermine PA as a critical regulator of mTOR complexes, while keeps the possibility that PA might be one of many players that ensures the stability of mTORC2.

Although future quantitative experiments can improve our model, it is a useful framework to understand the dynamics of signal integration and transduction by mTOR. While it is claimed that PA helps with stabilizing the mTORC2 more than mTORC1, our model reveals that the actual mechanism by which rapamycin affects mTORC2 integrity is through *Le Chatelier*’s principle. This finding has translational implications, and it will be a useful approach to design rapalogs that target one of the two mTOR complexes more selectively only by modifying their affinity towards either mTORC1 or the free mTOR.

## Funding Statement

The authors received no specific funding for this work.

## References

1. Tavassoly I: Dynamics of Cell Fate Decision Mediated by the Interplay of Autophagy and Apoptosis in Cancer Cells: Mathematical Modeling and Experimental Observations: Springer; 2015.

2. Hanahan D, Weinberg RA: Hallmarks of cancer: the next generation. cell 2011, 144(5):646–674.

3. Kreeger PK, Lauffenburger DA: Cancer systems biology: a network modeling perspective. Carcinogenesis 2009, 31(1):2–8.

4. Tavassoly I, Parmar J, Shajahan-Haq A, Clarke R, Baumann W, Tyson J: Dynamic modeling of the interaction between autophagy and apoptosis in mammalian cells. CPT: pharmacometrics & systems pharmacology 2015, 4(4):263–272.

5. Tavassoly I, Goldfarb J, Iyengar R: Systems biology primer: the basic methods and approaches. Essays in biochemistry 2018, 62(4):487–500.

6. Sorribes I, Basu A, Brady R, Enriquez-Navas P, Feng X, Kather J, Nerlakanti N, Stephens R, Strobl M, Tavassoly I: Harnessing patient-specific response dynamics to optimize evolutionary therapies for metastatic clear cell renal cell carcinoma-Learning to adapt. bioRxiv 2019:563130.

7. Tavassoly I, Hu Y, Zhao S, Mariottini C, Boran A, Chen Y, Li L, Tolentino RE, Jayaraman G, Goldfarb J et al: Systems therapeutics analyses identify genomic signatures defining responsiveness to allopurinol and combination therapy for lung cancer. bioRxiv 2018.

8. Hornberg JJ, Bruggeman FJ, Westerhoff HV, Lankelma J: Cancer: a systems biology disease. Biosystems 2006, 83(2-3):81–90.

9. Tyson JJ, Baumann WT, Chen C, Verdugo A, Tavassoly I, Wang Y, Weiner LM, Clarke R: Dynamic modelling of oestrogen signalling and cell fate in breast cancer cells. Nature Reviews Cancer 2011, 11(7):523.

10. Laplante M, Sabatini DM: mTOR signaling at a glance. Journal of cell science 2009, 122(20):3589–3594.

11. Saxton RA, Sabatini DM: mTOR signaling in growth, metabolism, and disease. Cell 2017, 168(6):960–976.

12. Ginzberg MB, Chang N, D’Souza H, Patel N, Kafri R, Kirschner MW: Cell size sensing in animal cells coordinates anabolic growth rates and cell cycle progression to maintain cell size uniformity. Elife 2018, 7:e26957.

13. Clarke R, Cook KL, Hu R, Facey CO, Tavassoly I, Schwartz JL, Baumann WT, Tyson JJ, Xuan J, Wang Y: Endoplasmic reticulum stress, the unfolded protein response, autophagy, and the integrated regulation of breast cancer cell fate. Cancer research 2012, 72(6):1321–1331.

14. Caron E, Ghosh S, Matsuoka Y, Ashton-Beaucage D, Therrien M, Lemieux S, Perreault C, Roux PP, Kitano H: A comprehensive map of the mTOR signaling network. Molecular systems biology 2010, 6(1):453.

15. Wang G, Krueger GR: Computational analysis of mTOR signaling pathway: Bifurcation, carcinogenesis, and drug discovery. Anticancer research 2010, 30(7):2683–2688.

16. Kim Y, Powathil G, Kang H, Trucu D, Kim H, Lawler S, Chaplain M: Strategies of eradicating glioma cells: a multi-scale mathematical model with MiR-451-AMPK-mTOR control. PloS one 2015, 10(1):e0114370.

17. Kapuy O, Vinod P, Bánhegyi G: mTOR inhibition increases cell viability via autophagy induction during endoplasmic reticulum stress–An experimental and modeling study. FEBS open bio 2014, 4(1):704–713.

18. Laplante M, Sabatini DM: mTOR signaling in growth control and disease. Cell 2012, 149(2):274–293.

19. Hay N, Sonenberg N: Upstream and downstream of mTOR. Genes & development 2004, 18(16):1926–1945.

20. Meng D, Frank AR, Jewell JL: mTOR signaling in stem and progenitor cells. Development 2018, 145(1):dev152595.

21. Guertin DA, Sabatini DM: The pharmacology of mTOR inhibition. Sci Signal 2009, 2(67):pe24–pe24.

22. Alessi DR, Pearce LR, García-Martínez JM: New insights into mTOR signaling: mTORC2 and beyond. Sci Signal 2009, 2(67):pe27–pe27.

23. Abraham RT, Wiederrecht GJ: Immunopharmacology of rapamycin. Annual review of immunology 1996, 14(1):483–510.

24. Sarbassov DD, Ali SM, Sengupta S, Sheen J-H, Hsu PP, Bagley AF, Markhard AL, Sabatini DM: Prolonged rapamycin treatment inhibits mTORC2 assembly and Akt/PKB. Molecular cell 2006, 22(2):159–168.

25. Demidenko ZN, Zubova SG, Bukreeva EI, Pospelov VA, Pospelova TV, Blagosklonny MV: Rapamycin decelerates cellular senescence. Cell cycle 2009, 8(12):1888–1895.

26. Dancey JE: Clinical development of mammalian target of rapamycin inhibitors. Hematology/Oncology Clinics 2002, 16(5):1101–1114.

27. Kahan BD, Camardo JS: Rapamycin: Clinical Results and Future Opportunities1. Transplantation 2001, 72(7):1181–1193.

28. Lamming DW, Ye L, Sabatini DM, Baur JA: Rapalogs and mTOR inhibitors as anti-aging therapeutics. The Journal of clinical investigation 2013, 123(3):980–989.

29. Fang Y, Westbrook R, Hill C, Boparai RK, Arum O, Spong A, Wang F, Javors MA, Chen J, Sun LY: Duration of rapamycin treatment has differential effects on metabolism in mice. Cell metabolism 2013, 17(3):456–462.

30. Fang Y, Bartke A: Prolonged rapamycin treatment led to beneficial metabolic switch. Aging (Albany NY) 2013, 5(5):328.

31. Ye L, Varamini B, Lamming DW, Sabatini DM, Baur JA: Rapamycin has a biphasic effect on insulin sensitivity in C2C12 myotubes due to sequential disruption of mTORC1 and mTORC2. Frontiers in genetics 2012, 3:177.

32. Tyson JJ, Chen K, Novak B: Network dynamics and cell physiology. Nature reviews Molecular cell biology 2001, 2(12):908.

33. Tyson JJ, Novák B: Functional motifs in biochemical reaction networks. Annual review of physical chemistry 2010, 61:219–240.

34. Atkins P, De Paula J, Keeler J: Atkins’ physical chemistry: Oxford university press; 2018.

35. Fang Y, Vilella-Bach M, Bachmann R, Flanigan A, Chen J: Phosphatidic acid-mediated mitogenic activation of mTOR signaling. Science 2001, 294(5548):1942–1945.

36. Foster DA: Regulation of mTOR by phosphatidic acid? Cancer research 2007, 67(1):1–4.

37. Ferron GM, Conway WD, Jusko WJ: Lipophilic benzamide and anilide derivatives as high-performance liquid chromatography internal standards: application to sirolimus (rapamycin) determination. Journal of Chromatography B: Biomedical Sciences and Applications 1997, 703

38. Vinod PKU, Venkatesh KV: Quantification of the effect of amino acids on an integrated mTOR and insulin signaling pathway. Molecular BioSystems 2009, 5(10):1163-1173.

39. Banaszynski LA, Liu CW, Wandless TJ: Characterization of the FKBP⊙ Rapamycin⊙ FRB Ternary Complex. Journal of the American Chemical Society 2005, 127(13):4715-4721.

40. Geiger T, Wehner A, Schaab C, Cox J, Mann M: Comparative proteomic analysis of eleven common cell lines reveals ubiquitous but varying expression of most proteins. Molecular & Cellular Proteomics 2012:mcp. M111. 014050.

41. MacDonald A, Scarola J, Burke JT, Zimmerman JJ: Clinical pharmacokinetics and therapeutic drug monitoring of sirolimus. Clinical therapeutics 2000, 22:B101-B121.

42. Albawardi A, Almarzooqi S, Saraswathiamma D, Abdul-Kader HM, Souid A-K, Alfazari AS: The mTOR inhibitor sirolimus suppresses renal, hepatic, and cardiac tissue cellular respiration. International journal of physiology, pathophysiology and pharmacology 2015, 7(1):54.

43. Chakrabarti S, Bhattacharya S, Bhattacharya SK: Biochemical engineering: cues from cells. TRENDS in Biotechnology 2003, 21(5):204-209.

44. Harrison DE, Strong R, Sharp ZD, Nelson JF, Astle CM, Flurkey K, Nadon NL, Wilkinson JE, Frenkel K, Carter CS: Rapamycin fed late in life extends lifespan in genetically heterogeneous mice. nature 2009, 460(7253):392.

45. Leontieva OV, Paszkiewicz GM, Blagosklonny MV: Mechanistic or mammalian target of rapamycin (mTOR) may determine robustness in young male mice at the cost of accelerated aging. Aging (Albany NY) 2012, 4(12):899.

46. Leontieva OV, Paszkiewicz GM, Blagosklonny MV: Weekly administration of rapamycin improves survival and biomarkers in obese male mice on high-fat diet. Aging cell 2014, 13(4):616-622.

47. Arriola Apelo SI, Neuman JC, Baar EL, Syed FA, Cummings NE, Brar HK, Pumper CP, Kimple ME, Lamming DW: Alternative rapamycin treatment regimens mitigate the impact of rapamycin on glucose homeostasis and the immune system. Aging cell 2016, 15(1):28-38.

48. Arriola Apelo SI, Pumper CP, Baar EL, Cummings NE, Lamming DW: Intermittent administration of rapamycin extends the life span of female C57BL/6J mice. Journals of Gerontology Series A: Biomedical Sciences and Medical Sciences 2016, 71(7):876-881.

49. Roulin D, Cerantola Y, Dormond-Meuwly A, Demartines N, Dormond O: Targeting mTORC2 inhibits colon cancer cell proliferation in vitro and tumor formation in vivo. Molecular cancer 2010, 9(1):57.

50. Lamanuzzi A, Saltarella I, Desantis V, Frassanito MA, Leone P, Racanelli V, Nico B, Ribatti D, Ditonno P, Prete MJO: Inhibition of mTOR complex 2 restrains tumor angiogenesis in multiple myeloma. 2018, 9(29):20563.

51. Farhan MA, Carmine-Simmen K, Lewis JD, Moore RB, Murray AGJPo: Endothelial cell mTOR complex-2 regulates sprouting angiogenesis. 2015, 10(8):e0135245.

52. Werfel TA, Wang S, Jackson MA, Kavanaugh TE, Joly MM, Lee LH, Hicks DJ, Sanchez V, Ericsson PG, Kilchrist KVJCr: Selective mTORC2 inhibitor therapeutically blocks breast cancer cell growth and survival. 2018, 78(7):1845–1858.

53. Zou Z, Chen J, Yang J, Bai XJCcdt: Targeted inhibition of Rictor/mTORC2 in cancer treatment: a new era after rapamycin. 2016, 16(4):288–304.

54. Guba M, von Breitenbuch P, Steinbauer M, Koehl G, Flegel S, Hornung M, Bruns CJ, Zuelke C, Farkas S, Anthuber M: Rapamycin inhibits primary and metastatic tumor growth by antiangiogenesis: involvement of vascular endothelial growth factor. Nature medicine 2002, 8(2):128.

55. Koehl G, Guba M, Seeliger H, Steinbauer M, Anthuber M, Jauch K-W, Geissler E: Rapamycin treatment at immunosuppressive doses affects tumor blood vessel circulation. In: Transplantation proceedings: 2003. 2135–2136.

56. Mannick JB, Morris M, Hockey H-UP, Roma G, Beibel M, Kulmatycki K, Watkins M, Shavlakadze T, Zhou W, Quinn D: TORC1 inhibition enhances immune function and reduces infections in the elderly. Science translational medicine 2018, 10(449):eaaq1564.

57. Mannick JB, Del Giudice G, Lattanzi M, Valiante NM, Praestgaard J, Huang B, Lonetto MA, Maecker HT, Kovarik J, Carson S: mTOR inhibition improves immune function in the elderly. Science translational medicine 2014, 6(268):268ra179–268ra179.

58. Toschi A, Lee E, Gadir N, Ohh M, Foster DA: Differential dependence of hypoxia-inducible factors 1α and 2α on mTORC1 and mTORC2. Journal of Biological Chemistry 2008, 283(50):34495–34499.

