## Supplementary Materials for "Dynamic Modeling of Signal Transduction by mTOR Complexes in Cancer"

**Supplementary Table S1**: Kinetic properties of rapamycin based on Ferron et al. ^1^: K_abs_, A, B, 𝛂 and 𝛃 are extracted from Ferron et al. ^1^; K_el_ is calculated based on these values using the equation presented in the table.

| Parameter | Amounts | Units | Descriptions |
| --- | --- | --- | --- |
| K_abs_ | 2.77 | 1/h | Absorption rate constant |
| A | 7.89 | ng/ml | Distribution phase intercept |
| α | 0.28 | 1/h | Distribution phase slope |
| B | 1.06 | ng/ml | Elimination phase intercept |
| 𝛽 | 0.011 | 1/h | Elimination phase slope |
| K_el_ | 0.0718632 | 1/h | Elimination rate constant calculated by $K_{el}=\frac{\alpha.\beta.(A+B)}{A.\beta+B.\alpha}$ |

**Supplementary Table S2**: Model specifications extracted from literature: Vinod et al. ^2^ assumed k_c1f_ to be equal to 1 (nM.min)^-1^ and calculated k_c1d_ based on their model. Banaszynski et al. ^3^ calculated k_i1f_ and k_i1d_ as the association and dissociation rate constants of binding of Rapamycin-FKBP12 to FRB domain of mTOR. Geiger et al.^4^ measured the copy number of the proteome of 11 cell lines including HEK293 and reported the amount of mTOR, Raptor, and Rictor as molecule per cell. The particle number is transformed to the number of moles by Avogadro number, and the molarity of each species is calculated by assuming the volume of HEK293 equal to 5.0E-12 liter. k_c1f_= formation rate constant of mTORC1; k_c1d_= dissociation rate constant of mTORC1; k_i1f_= association rate constant of Rapamycin and mTOR; k_i1r_= dissociation (reverse reaction) rate constant of Rapamycin-mTOR complex; (nM.min)^-1^= per nanoMolar per minute; min^-1^= per minute; (M.S)^-1^= per Molar per Second; S^-1^= per Second.

| Reference | mTORC1 | | mTOR-Rapamycin | | Concentrations | | |
| --- | --- | --- | --- | --- | --- | --- | --- |
|  | k_C1f_ | k_C1d_ | k_i1f_ | K_i1r_ | [mTOR] | [Raptor] | [Rictor] |
| Geiger et al. 2012 |  |  |  |  | 1131000 Molecule  or  1.878114*10^-18^ mole  or  3.756228*10^-7^ Molar | 964000 Molecule  or  1.600797*10^-18^ mole  or  3.201594*10^-7^ Molar | 266000 Molecule  or  4.417137*10^-19^ mole  or  8.834274*10^-8^ Molar |
| Vinod et al. 2009 | 1 (nM.min)^-1^  or  1.6666666*10^7^  (M.S)^-1^ | 5 min^-1^  or  8.333*10^-2^  S^-1^ |  |  |  |  |  |
| Banaszynski et al. 2004 |  |  | 1.92*10^6^  (M.S)^-1^ | 2.2*10^-2^ S^-1^ |  |  |  |

**Supplementary Table S3**: Minimum and maximum residual active concentration and percentage of mTORC1 and mTORC2 for each schedule, scanned in ± 20% dose area, was calculated. [mTORC1] is the residual mTORC1 concentration, mTORC1% is the residual active percent of mTORC1, [mTORC2] is the residual concentration of mTORC2, and mTORC2% is the residual active percent of mTORC2.

|  |  |  | mTORC1 | |  | mTORC2 | |
| --- | --- | --- | --- | --- | --- | --- | --- |
|  |  |  | [mTORC1] | mTORC1% |  | [mTORC2] | mTORC2% |
| Drug-free period |  |  | 2.7217*10^-7^ | 100% |  | 7.51*10^-8^ | 100% |
| 8.0*10^-20^ mole/Day | 80% | Min | 1.7123*10^-7^ | 62.913 |  | 6.4167*10^-8^ | 85.442 |
|  |  | Avg. |  | 64.4965 |  |  | 86.2055 |
|  |  | Max | 1.7985*10^-7^ | 66.080 |  | 6.5314*10^-8^ | 86.969 |
|  | 90% | Min | 1.6329*10^-7^ | 59.996 |  | 6.3045*10^-8^ | 83.948 |
|  |  | Avg. |  | 61.763 |  |  | 84.8335 |
|  |  | Max | 1.7291*10^-7^ | 63.530 |  | 6.4375*10^-8^ | 85.719 |
|  | 100% | Min | 1.56*10^-7^ | 57.317 |  | 6.1968*10^-8^ | 82.514 |
|  |  | Avg. |  | 59.2605 |  |  | 83.5255 |
|  |  | Max | 1.6658*10^-7^ | 61.204 |  | 6.3487*10^-8^ | 84.537 |
|  | 110% | Min | 1.4924*10^-7^ | 54.833 |  | 6.0933*10^-8^ | 81.136 |
|  |  | Avg. |  | 56.953 |  |  | 82.2765 |
|  |  | Max | 1.6078*10^-7^ | 59.073 |  | 6.2646*10^-8^ | 83.417 |
|  | 120% | Min | 1.4298*10^-7^ | 52.533 |  | 5.9936*10^-8^ | 79.808 |
|  |  | Avg. |  | 54.822 |  |  | 81.0805 |
|  |  | Max | 1.5544*10^-7^ | 57.111 |  | 6.1847*10^-8^ | 82.353 |
| 5.6*10^-19^ mole/Day | 80% | Min | 4.2297*10^-8^ | 15.541 |  | 3.6405*10^-8^ | 48.475 |
|  |  | Avg. |  | 21.4465 |  |  | 54.745 |
|  |  | Max | 7.4443*10^-8^ | 27.352 |  | 4.5822*10^-8^ | 61.015 |
|  | 90% | Min | 3.7239*10^-8^ | 13.682 |  | 3.4576*10^-8^ | 46.040 |
|  |  | Avg. |  | 19.7115 |  |  | 52.7555 |
|  |  | Max | 7.006*10^-8^ | 25.741 |  | 4.4663*10^-8^ | 59.471 |
|  | 100% | Min | 3.3156*10^-8^ | 12.182 |  | 3.3008*10^-8^ | 43.952 |
|  |  | Avg. |  | 18.2765 |  |  | 51.0335 |
|  |  | Max | 6.6331*10^-8^ | 24.371 |  | 4.3644*10^-8^ | 58.115 |
|  | 110% | Min | 2.9813*10^-8^ | 10.954 |  | 3.1653*10^-8^ | 42.148 |
|  |  | Avg. |  | 17.0695 |  |  | 49.526 |
|  |  | Max | 6.3103*10^-8^ | 23.185 |  | 4.2735*10^-8^ | 56.904 |
|  | 120% | Min | 2.7034*10^-8^ | 9.933 |  | 3.0469*10^-8^ | 40.571 |
|  |  | Avg. |  | 16.037 |  |  | 48.191 |
|  |  | Max | 6.0262*10^-8^ | 22.141 |  | 4.1914*10^-8^ | 55.811 |
| 5.6*10^-19^ mole/Week | 80% | Min | 1.4077*10^-7^ | 51.721 |  | 5.9647*10^-8^ | 79.423 |
|  |  | Avg. |  | 63.442 |  |  | 85.204 |
|  |  | Max | 2.0457*10^-7^ | 75.163 |  | 6.833*10^-8^ | 90.985 |
|  | 90% | Min | 1.2948*10^-7^ | 47.573 |  | 5.7743*10^-8^ | 76.888 |
|  |  | Avg. |  | 60.6035 |  |  | 83.602 |
|  |  | Max | 2.0041*10^-7^ | 73.634 |  | 6.7827*10^-8^ | 90.316 |
|  | 100% | Min | 1.1899*10^-7^ | 43.719 |  | 5.5823*10^-8^ | 74.332 |
|  |  | Avg. |  | 58.008 |  |  | 82.023 |
|  |  | Max | 1.9677*10^-7^ | 72.297 |  | 6.7375*10^-8^ | 89.714 |
|  | 110% | Min | 1.0925*10^-7^ | 40.140 |  | 5.3983*10^-8^ | 71.881 |
|  |  | Avg. |  | 55.6325 |  |  | 80.5285 |
|  |  | Max | 1.9358*10^-7^ | 71.125 |  | 6.6971*10^-8^ | 89.176 |
|  | 120% | Min | 1.0023*10^-7^ | 36.826 |  | 5.2138*10^-8^ | 69.425 |
|  |  | Avg. |  | 53.459 |  |  | 79.0595 |
|  |  | Max | 1.9077*10^-7^ | 70.092 |  | 6.6609*10^-8^ | 88.694 |
| 2.24*10^-18^ mole/Week | 80% | Min | 2.1655*10^-8^ | 7.956 |  | 2.8217*10^-8^ | 37.573 |
|  |  | Avg. |  | 35.124 |  |  | 61.1355 |
|  |  | Max | 1.6954*10^-7^ | 62.292 |  | 6.3608*10^-8^ | 84.698 |
|  | 90% | Min | 1.765*10^-8^ | 6.485 |  | 2.6179*10^-8^ | 34.859 |
|  |  | Avg. |  | 34.1165 |  |  | 59.626 |
|  |  | Max | 1.6806*10^-7^ | 61.748 |  | 6.3379*10^-8^ | 84.393 |
|  | 100% | Min | 1.4786*10^-8^ | 5.433 |  | 2.4572*10^-8^ | 32.719 |
|  |  | Avg. |  | 33.3665 |  |  | 58.4295 |
|  |  | Max | 1.6684*10^-7^ | 61.300 |  | 6.3189*10^-8^ | 84.140 |
|  | 110% | Min | 1.2659*10^-8^ | 4.651 |  | 2.3269*10^-8^ | 30.984 |
|  |  | Avg. |  | 32.7825 |  |  | 57.4535 |
|  |  | Max | 1.6579*10^-7^ | 60.914 |  | 6.3026*10^-8^ | 83.923 |
|  | 120% | Min | 1.1029*10^-8^ | 4.052 |  | 2.2184*10^-8^ | 29.539 |
|  |  | Avg. |  | 32.316 |  |  | 56.6355 |
|  |  | Max | 1.6488*10^-7^ | 60.580 |  | 6.2883*10^-8^ | 83.732 |

**Supplementary Table S4**: Average concentration and average residual active percentage of mTORC1 and mTORC2 in different absorption and elimination rate constants (macro-constants) of rapamycin: K_abs_ and K_el_ were scanned in an area of ± 2logs with three steps for each. The input dose is 5.6*10^-19^ mole/Day (regimen Ⅱ). The percentages are rounded to two decimal point. Amounts corresponding to the initial values of parameters are boldfaced. [mTORC1] is the residual mTORC1 concentration, mTORC1% is the residual active percent of mTORC1, [mTORC2] is the residual concentration of mTORC2, mTORC2% is the residual active percent of mTORC2, k_abs_ is the absorption rate constant, and K_el_ is the elimination rate constant.

|  |  | mTORC1 | |  | mTORC2 | |
| --- | --- | --- | --- | --- | --- | --- |
|  |  | [mTORC1]_avg._ | mTORC1%_avg._ |  | [mTORC2]_avg._ | mTORC2%_avg._ |
|  | Drug-free period | 2.7217*10^-7^ | 100% |  | 7.51*10^-8^ | 100% |
| K_abs_ = 0.0277 | K_el_ = 0.000718632 | 4.1106*10^-10^ | 0.15 |  | 6.4689*10^-9^ | 8.61 |
|  | K_el_ = 0.0718632 | 4.7521*10^-8^ | 17.46 |  | 3.8089*10^-8^ | 50.72 |
|  | K_el_ = 7.18632 | 2.6011*10^-7^ | 95.57 |  | 7.4059*10^-8^ | 98.61 |
| K_abs_ = 2.77 | K_el_ = 0.000718632 | 4.0726*10^-10^ | 0.15 |  | 6.4408*10^-9^ | 8.58 |
|  | K_el_ = 0.0718632 | **4.9746*10^-8^** | **18.28** |  | **3.8327*10^-8^** | **51.04** |
|  | K_el_ = 7.18632 | 2.4072*10^-7^ | 88.45 |  | 7.2037*10^-8^ | 95.92 |
| K_abs_ = 277 | K_el_ = 0.000718632 | 4.0703*10^-10^ | 0.15 |  | 6.4369*10^-9^ | 8.57 |
|  | K_el_ = 0.0718632 | 4.9069*10^-8^ | 18.03 |  | 3.8041*10^-8^ | 50.65 |
|  | K_el_ = 7.18632 | 2.2998*10^-7^ | 84.50 |  | 7.0884*10^-8^ | 94.39 |

**Supplementary Table S5**: Average percent of remaining active mTORC1 and mTORC2 in the parameter scan for association and dissociation rate constants namely k_i1f_, k_i1r_, k_i2f_, k_i2r_, k_i3f_, and k_i3r_. Every rate constant was scanned separately in an area of ± 3logs in 7 steps. The reference dose is 5.6*10^-19^ mole/Day (regimen Ⅱ). k_i1f_ is the rapamycin binding to mTOR rate constant, k_i1r_ is the rapamycin-mTOR complex dissociation rate constant, k_i2f_ is the rapamycin binding to mTORC1 rate constant, k_i2r_ is the releasing rapamycin from mTORC1-Rapamycin rate constant, k_i3f_ is the releasing Raptor from mTORC1-Rapamycin complex rate constant, k_i3r_ is the rate constant of association of Raptor to mTOR-Rapamycin complex, mTORC1%_avg._ is the average residual active percentage of mTORC1 and mTORC2%_avg._ is the average residual active percentage of mTORC2.

|  |  | mTORC1%_avg._ | mTORC2%_avg._ |
| --- | --- | --- | --- |
| k_i1f_ = 1.92*10^6^ | -3Logs | 19.238 | 56.908 |
|  | -2Logs | 19.227 | 56.840 |
|  | -Log | 19.124 | 56.181 |
|  | 1 | 18.276 | 51.036 |
|  | Log | 14.324 | 32.251 |
|  | 2Logs | 6.595 | 10.791 |
|  | 3Logs | 1.421 | 1.898 |
| k_i1r_ = 2.2*10^-2^ | -3Logs | 0.113 | 0.147 |
|  | -2Logs | 1.030 | 1.386 |
|  | -Log | 6.617 | 10.898 |
|  | 1 | 18.277 | 51.035 |
|  | Log | 22.499 | 82.517 |
|  | 2Logs | 23.090 | 88.390 |
|  | 3Logs | 23.150 | 89.056 |
| k_i2f_ = 1.92*10^6^ | -3Logs | 83.440 | 83.682 |
|  | -2Logs | 80.560 | 82.875 |
|  | -Log | 60.256 | 76.031 |
|  | 1 | 18.277 | 51.033 |
|  | Log | 0.823 | 33.539 |
|  | 2Logs | 0.334 | 29.704 |
|  | 3Logs | 0.034 | 29.206 |
| k_i2r_ = 2.2*10^-2^ | -3Logs | 7.380 | 39.621 |
|  | -2Logs | 7.499 | 39.766 |
|  | -Log | 8.661 | 41.157 |
|  | 1 | 18.277 | 51.033 |
|  | Log | 54.484 | 73.614 |
|  | 2Logs | 79.208 | 82.477 |
|  | 3Logs | 83.294 | 83.639 |
| k_i3f_ = 10^-2^ | -3Logs | 16.829 | 64.233 |
|  | -2Logs | 16.853 | 64.035 |
|  | -Log | 17.082 | 62.149 |
|  | 1 | 18.278 | 51.033 |
|  | Log | 19.314 | 40.093 |
|  | 2Logs | 19.485 | 38.224 |
|  | 3Logs | 19.503 | 38.025 |
| k_i3r_ = 1.0*10^-5^ | -3Logs | 18.277 | 51.031 |
|  | -2Logs | 18.277 | 51.031 |
|  | -Log | 18.277 | 51.031 |
|  | 1 | 18.277 | 51.031 |
|  | Log | 18.277 | 51.031 |
|  | 2Logs | 18.277 | 51.031 |
|  | 3Logs | 18.277 | 51.031 |

**Supplementary Figure S1**: Temporal concentrations of mTORC1 and mTORC2 for different values of association and dissociation rate constants of all species namely kc1f, kc1d, kc2f, kc2d, ki1f, ki1r, ki2f, ki2r, ki3f and ki3r. Every rate constant was scanned separately in an area of ± 3logs in 7 steps. The reference dose of Rapamycin is 5.6*10-19 mole/Day (regimen II). kc1f = mTORC1 formation rate constant; kc1d = mTORC1 dissociation rate constant; kc2f = mTORC2 formation rate constant; kc2d = mTORC2 dissociation rate constant; ki1f = Rapamycin binding to mTOR rate constant; ki1r = Rapamycin-mTOR complex dissociation rate constant; ki2f = Rapamycin binding to mTORC1 rate constant; ki2r = Releasing Rapamycin from mTORC1-Rapamycin rate constant; ki3f = Releasing Raptor from mTORC1-Rapamycin complex rate constant; ki3r = rate constant of association of Raptor to mTOR-Rapamycin complex.

**REFERENCES**

1.	Ferron, G.M., Conway, W.D. & Jusko, W.J. Lipophilic benzamide and anilide derivatives as high-performance liquid chromatography internal standards: application to sirolimus (rapamycin) determination. *Journal of Chromatography B: Biomedical Sciences and Applications* **703**, 243-251 (1997).

2.	Vinod, P.K.U. & Venkatesh, K.V. Quantification of the effect of amino acids on an integrated mTOR and insulin signaling pathway. *Molecular BioSystems* **5**, 1163-1173 (2009).

3.	Banaszynski, L.A., Liu, C.W. & Wandless, T.J. Characterization of the FKBP⊙ Rapamycin⊙ FRB Ternary Complex. *Journal of the American Chemical Society* **127**, 4715-4721 (2005).

4.	Geiger, T., Wehner, A., Schaab, C., Cox, J. & Mann, M. Comparative proteomic analysis of eleven common cell lines reveals ubiquitous but varying expression of most proteins. *Molecular & Cellular Proteomics*, mcp. M111. 014050 (2012).
