## Supplementary figures and images for "Dynamic Modeling of Signal Transduction by mTOR Complexes in Cancer"

### Supplementary Figure 1

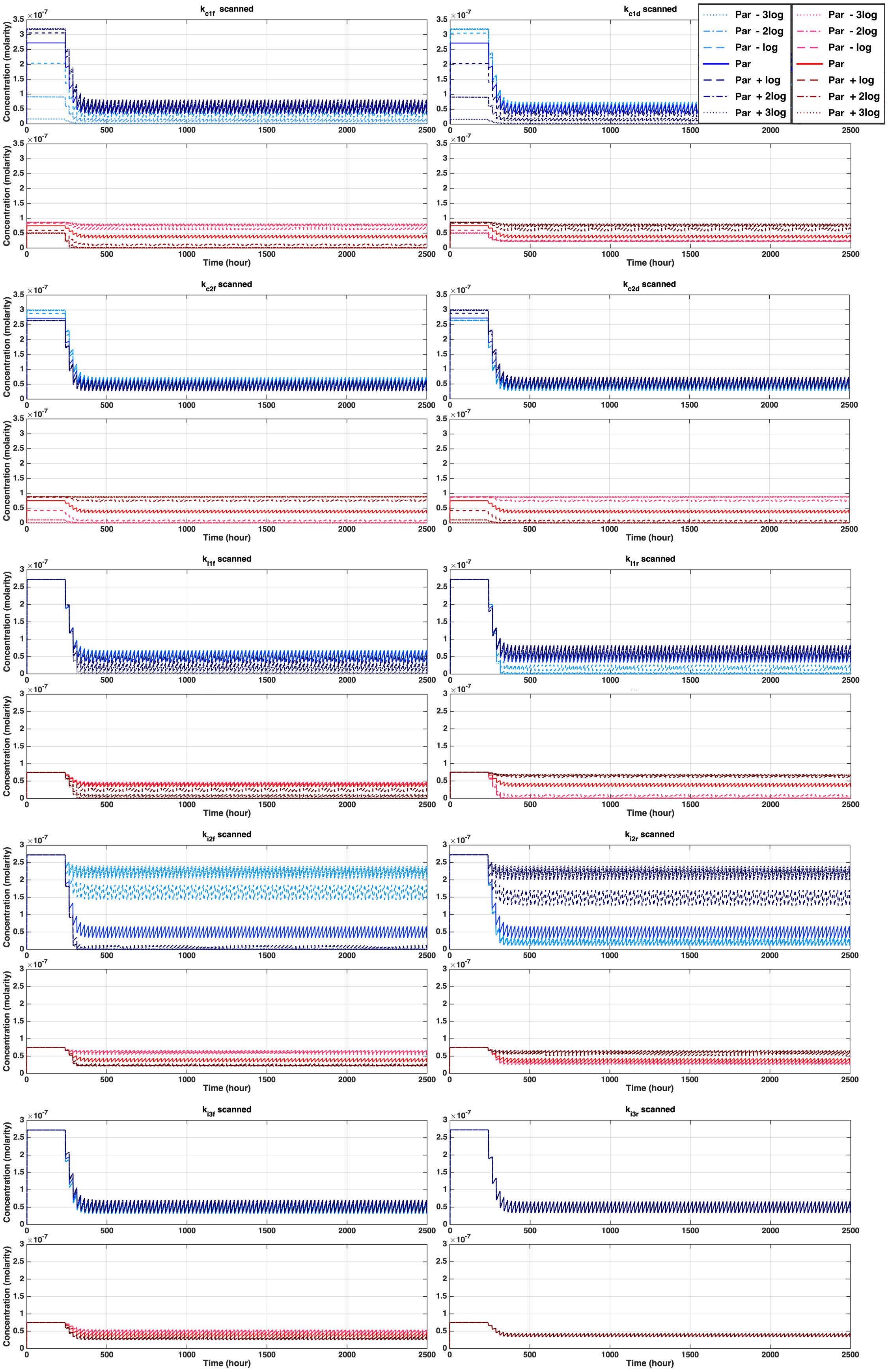
